# Tissue ingrowth markedly reduces mechanical anisotropy and stiffness in fibre direction of highly aligned electrospun polyurethane scaffolds

**DOI:** 10.1101/779942

**Authors:** Hugo Krynauw, Jannik Buescher, Josepha Koehne, Loes Verrijt, Georges Limbert, Neil H Davies, Deon Bezuidenhout, Thomas Franz

**Author notes:** Corresponding author: Thomas Franz, PhD, Division of Biomedical Engineering, Department of Human Biology, Faculty of Health Sciences, University of Cape Town, Private Bag X3, Observatory 7935, South Africa.

## Abstract

**Purpose:** The lack of long-term patency of synthetic vascular grafts currently available on the market has directed research towards improving the performance of small diameter grafts. Improved radial compliance matching and tissue ingrowth into the graft scaffold are amongst the main goals for an ideal vascular graft.

**Methods:** Biostable polyurethane scaffolds were manufactured by electrospinning and implanted in subcutaneous and circulatory positions in the rat for 7, 14 and 28 days. Scaffold morphology, tissue ingrowth, and mechanical properties of the scaffolds were assessed before implantation and after retrieval.

**Results:** Tissue ingrowth after 24 days was 96.5 ± 2.3% in the subcutaneous implants and 77.8 ± 5.4% in the circulatory implants. Over the 24 days implantation, the elastic modulus at 12% strain decreased by 59% in direction of the fibre alignment whereas it increased by 1379% transverse to the fibre alignment of the highly aligned scaffold of the subcutaneous implants. The lesser aligned scaffold of the circulatory graft implants exhibited an increase of the elastic modulus at 12% strain by 77% in circumferential direction.

**Conclusion:** Based on the observations, it is proposed that the mechanism underlying the softening of the highly aligned scaffold in the predominant fibre direction is associated with scaffold compaction and local displacement of fibres by the newly formed tissue. The stiffening of the scaffold, observed transverse to highly aligned fibres and for more a random fibre distribution, represents the actual mechanical contribution of the tissue that developed in the scaffold.

## 1. Introduction

Regenerative medicine has emerged as one of the most dynamic drivers in the development of advanced engineered biomaterial solutions for tissue engineering applications.^5, 24^ One crucial element in regenerative medicine is scaffolds that facilitate and guide the engineering and regeneration of biological tissues according to the application.

The ideal scaffold facilitates tissue engineering and regeneration such that the new tissue constructs biologically and mechanically mimic the healthy host tissue.^5^ In cardiovascular regenerative therapies, one of the major challenges remains small-diameter vascular grafts.^29^ For these grafts, a key factor for the long term success is porosity.^29^ One method of introducing porosity in scaffolds for the vascular tissue engineering is electrospinning of polymers.^2, 3, 6, 7, 9, 10, 13-15, 17, 18, 22^

For electrospinning of polymeric networks, the process parameters allow controlling the degree of fibre alignment^1, 26^ which affects the mechanical properties of the fibrous scaffold. Scaffolds with fibres randomly distributed typically exhibit similar properties in different directions whereas the alignment of fibres predominantly in one direction introduces mechanical anisotropy.^1^ Electrospun scaffolds with a high degree of fibre alignment exhibit a higher elastic modulus, i.e. stiffness, in fibre direction and a lower elastic modulus perpendicular to the fibre direction. The directional mechanical properties of the scaffold can be utilised to tailor structural properties. They add, however, complexity to the design process which needs to be adequately addressed.

Another factor that influences the structural and mechanical properties of the scaffold is the ingrowth of cells and tissue - whether through in vitro tissue culture or after implantation. When optimising scaffold-based engineered tissue constructs, the structural contribution of the new tissue should be considered in the design of the original scaffold. The incorporation of tissue in the scaffold is a transient process. As such, a gradual change of the mechanical properties of the scaffold-tissue construct is expected until tissue ingrowth and maturation are complete. Biomechanical properties of scaffolds are predominantly obtained prior to tissue culturing or implantation^2,4, 14, 28^ whereas the changes in mechanical properties due to tissue ingrowth after implantation have received very little attention. The use of a biostable polyurethane as scaffold material facilitates the isolation of the mechanical effects of tissue ingrowth from those of scaffold degradation that typically occur concurrently with tissue ingrowth in the case of biodegradable scaffolds.

## 2. Materials and Methods

### 2.1 Scaffold material

Pellethane© 2363–80AE (Dow Chemicals, USA), a medical grade aromatic poly(ether urethane) with a hard segment consisting of 4,4-methylenebis(phenylisocyanate) and 1,4-butanediol and a soft segment of poly(tetramethylene oxide) glycol (Mn = 1000 g/mole)^20^ was used.

### 2.2 Electrospinning and sample preparation

A 15 wt% Pellethane© solution was obtained by dissolving 8 g of Pellethane© pellets in 45.3 g of tetrahydrofuran (THF, Sigma Aldrich, Steinheim, Germany) at 37°C for 8 hours. Using a custom-made rig, the Pellethane® solution was electrospun from a hypodermic needle (SE400B syringe pump, Fresenius, Bad Homburg, Germany) onto a rotating and bi-directionally translating tubular target (hypodermic tubing, Small Parts, Loganport, IN, USA). The spinning process parameters were, for subcutaneous and circulatory samples, respectively: solution flow rate of 6 and 4.8 mL/h, target outer diameter of 25.0 and 2.2 mm, target rotating speed of 4400 and 9600 rpm, target translational speed of 2.6 mm/min, electrostatic field of 13 and 15 kV, and source-target distance of 250 mm (Note: Parameters were the same if only one value is given).

After completion of the spinning process, the electrospun structure on the target hypotube was submersed in ethanol for 5 minutes, removed from the mandrel and dried under vacuum (Townson & Mercer Ltd, Stretford, England; room temperature, 90 min) and trimmed on either end to discard regions with decreasing wall thickness. For subcutaneous implants, tubular scaffolds were cut into 9 × 18 mm rectangular samples with the longer edge aligned with either the circumferential (C) or axial (A) tube direction. For circulatory implants, scaffold tubes were cut into 11 mm long pieces. Samples were sterilised by submersion in 70% ethanol (room temperature, 24 h) and dried under vacuum (Townson & Mercer, Stretford, England, room temperature, 24 h). Samples for circulatory implants additionally underwent standard ETO sterilisation (55°C, 60% relative humidity, 12 h).

### 2.3 Physical characterisation of scaffold before implantation

Physical characterisation included microscopic analysis of fibre diameter and alignment, measurement of scaffold wall thickness and porosity.

#### 2.3.1 Fibre diameter and alignment

Scanning electron microscope (SEM) images were obtained (Nova NanoSEM 230, FEI, Hillsboro, OR, US) of the internal and external surfaces of samples after sputter-coating with gold (Polaron SC7640, Quorum Technologies, East Grinstead, England). The fibre diameter was measured with Scion Image (Scion Corporation, Frederick, USA) on x750 SEM micrographs (n=3 samples, two images per image side, both sides of sample, ten measurements per image).

Fibre alignment was determined by processing x100 SEM images in Fiji (based on ImageJ 1.48)^19^ by means of Fourier component analysis in the Directionality plug-in (written by JeanYves Tinevez) as used by Woolley et al. ^25^ The plug-in examines the power spectrum of Fourier transforms of the images in a polar coordinate system, measuring the power in user specified angular increments. A Gaussian function is fitted to the highest peak from which dispersion is calculated as the standard deviation of the Gaussian. This dispersion factor was used as an indication of fibre alignment. Smaller values indicate greater alignment.

#### 2.3.2 Scaffold and sample dimensions

Width, length and wall thickness of scaffold samples were measured on images captured on a Leica DFC280 stereo microscope using Leica IM500 imaging software (Leica Microsystems GmbH, Wetzlar, Germany). Six thickness measurements were recorded on both length edges of each sample as well as six measurements for sample width. The sample mass was determined using a Mettler Toledo XS105S analytical balance (Mettler Toledo, Greifensee, Switzerland).

#### 2.3.3 Scaffold porosity

Scaffold porosity, *P*, was determined by hydrostatic weighing typically employed for density determination. The porosity of fibrous networks is described as volumetric ratio

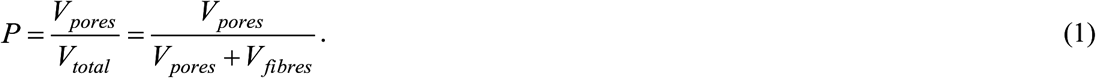

Expressing volumes as masses and considering that ethanol was used for hydrostatic weighing, providing *m*_*eth*.*pores*_ and *m*_*eth*.*total*_, Eq. 1 can be rewritten as

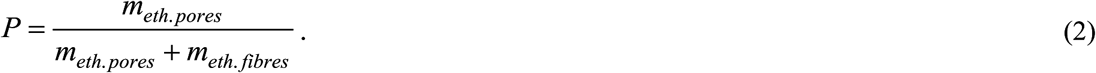

Introducing measurable quantities provides

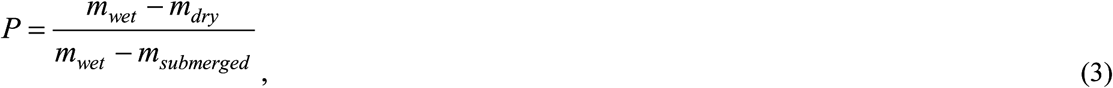

where *m*_*dry*_ is the mass of the dry scaffold, *m*_*submerged*_ is the mass of the scaffold submerged in ethanol, and *m*_*wet*_ is the mass of the scaffold after removal from the ethanol with ethanol retained in the pores. Equations 2 and 3 can be substituted as follows:

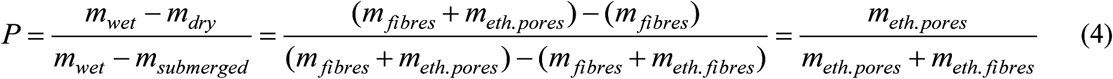

The porosity of each spun scaffold for each study was measured as described above with n = 1 per electrospun scaffold.

### 2.4 Study design

This study comprised subcutaneous and circulatory implants in a rat model with implant durations of 7, 14 and 28 days with n = 5 and 7 rats for subcutaneous and circulatory implant position, respectively, for each time point. For the subcutaneous implant position, each animal received five implants, four for mechanical testing and one for histological analysis. For the circulatory position, each animal received one graft implant.

### 2.5 Scaffold implantation

All animal experiments were authorised by the Faculty of Health Sciences Research Ethics Committee of the University of Cape Town and were performed in accordance with the National Institutes of Health (NIH, Bethesda, MD) guidelines.

Male Wistar albino rats with body mass 200-250 g for subcutaneous implants and 350-400 g for circulatory implants were used. Anaesthesia was induced by placing the animal in an inhalation chamber with an air flow of 5% isoflurane for 5 min. The animals were shaved and sterilized with iodine in the area of the incision. Anaesthesia was maintained by nose cone delivery of 1.5% isoflurane at an oxygen output of 1.5 L/min at 1 bar and 21°C. The animal’s body temperature was maintained throughout the surgical procedure by placing the animal on a custom-made heating pad at 37°C.

#### 2.5.1 Subcutaneous implants

Five longitudinal incisions of 1 cm were made, two on one and three on the other side of the dorsal midline, and a pocket of 2 cm in depth was blunt dissected subcutaneously at each incision. Scaffold samples were placed within the pockets and incisions closed with silk 4/0 (Ethicon Inc, Somerville, NJ) interrupted sutures in a subcuticular fashion.

#### 2.5.2 Circulatory graft implants

The aorta was exposed through a midline laparotomy and mobilised. Heparin was administered (1 mg/kg) prior to the cross-clamping of the aorta. The abdominal aorta was transected between the renal artery and the iliac bifurcation, the exposed aortic lumen was washed with heparin solution (1 mg/kg). The graft was anastomosed with interrupted sutures (9/0 Ethilon, Johnson & Johnson, New Brunswick, NJ) using an operation microscope (Zeiss Universal S3, Oberkochen, Germany). The distal clamp was carefully released to de-air the graft, then the proximal clamp was released. If there was no bleeding, the abdomen was closed in two layers with continuous sutures (2/0 Ethibond, Johnson & Johnson, New Brunswick, NJ). Buprenorphine (0,1mg/kg) was administered subcutaneously before abdomen was closed, as well as for the first 3 days post operatively twice daily.

### 2.6 Implant retrieval

Anaesthesia was induced by placing the animal in an inhalation chamber with an air flow of 5% isoflurane for 5 min.

#### 2.6.1 Subcutaneous implants

Rats were euthanized by inhalation of 5% halothane in air followed by a cardiac injection of 1 ml saturated KCl solution. The samples with surrounding tissue were excised.

#### 2.6.2 Circulatory graft implants

The abdominal area was shaved. Anaesthesia was maintained by nose cone delivery of 1.5% Isoflurane at an oxygen output of 1.5 l/min at 1 bar and 21°C. The graft implant site was exposed through a midline incision and dissected from adhesions. 1 mg/kg IV heparin was given via the inferior vena cava and allowed to circulate for 3 mins. The animal was then exsanguinated. After apnoea the chest was opened via sternotomy. A 20G cannula was inserted through the apex of the heart into the aorta. The aorta then was flushed with 200 ml NaCl 0.9% until clear fluid drained out of the opened right atrium. Perfusion fixation was performed with 100 ml paraformaldehyde. Thereafter the graft was excised and dissected in sections for mechanical testing and histological analysis.

#### 2.6.3 Processing of retrieved samples

Samples for mechanical testing were submerged in phosphate buffered saline solution (PBS) and stored at 4°C. Samples for histological analysis underwent tissue fixation in 10% formalin (Sigma Aldrich, Steinheim, Germany) for 24 hours, after which they were stored in a 70% ethanol solution.

### 2.7 Mechanical testing

Prior to testing, the thickness, length and width of retrieved subcutaneous samples were determined with a calliper. For retrieved graft samples, the wall thickness and width were measured using a stereo microscope (Leica DFC280 and Leica IM500 imaging software, Leica Microsystems GmbH, Wetzlar, Germany) and calliper, respectively.

Tensile testing was performed within 12 hours of implant retrieval with samples submerged in PBS at 37°C (Instron 5544, 10 N load cell; Instron, Norwood, USA). Flat subcutaneous samples were fixed using custom made clamps resulting in a gauge length of approximately 10 mm. The direction of tensile load represented either circumferential (C) or axial direction (A) of the original tubular scaffold. Circulatory graft samples were placed over two custom pin holders (hypodermic needles, outer diameter 0.9 mm) and loaded in circumferential direction. The loading protocol comprised one initial extension to 12% strain, five cycles between 12% and 8% strain, and a final extension to failure or the load limit of the load cell of 7.4 N (whichever came first) for subcutaneous samples, and to 17% strain or the load limit of the load cell (whichever came first) for circulatory graft samples (crosshead speed: 20 mm/min).

### 2.8 Histology

After fixation (4% paraformaldehyde, 24 h), samples underwent tissue processing and paraffin embedding. Sections of 3 μm thickness were cut from mid sample regions and stained with haematoxylin and eosin (H&E) for nuclei and cytoplasm. Images were acquired with a Nikon Eclipse 90i microscope with digital camera DXM-1200C (Nikon Corporation, Tokyo, Japan). Image analysis was performed to quantify tissue ingrowth by classifying area in the images as open space or tissue. VIS Visiopharm Integrator System (Visiopharm, Hørsholm, Denmark) was used to select a region of interest (ROI), segment the ROI into tissue and open space and then to calculate tissue ingrowth percentage per slide. Full segmentation involved three phases: 1) pre-processing the image; 2) segmenting by means of an untrained k-means clustering technique into nuclei, cytoplasm and extracellular matrix, and open space; and 3) post-processing to group all tissue and to remove small artefacts. Electrospun fibres did not stain and were thus counted as open space. To correct for this, the total area of each image was adjusted by the scaffold porosity.

### 2.9 Statistical analysis

Data were assessed for normality using Shapiro-Wilk test. One-way ANOVA was performed when more than two groups were compared by using Tukey HSD post-hoc analysis with p < 0.05 indicating statistical significance. Data are provided as mean ± standard deviation.

## 3. Results

### 3.1 Pre-implant scaffold morphology

**Figure 1** provides SEM micrographs of the electrospun scaffold for subcutaneous and circulatory implantation. Wall thickness, porosity, fibre diameter and fibre alignment of the scaffold for subcutaneous and graft implantation are summarised in **Table 1**. The scaffold for subcutaneous implantation featured generally long fibres with mostly uniform thickness that were highly aligned. Fairly low degree of fibre merging was observed, as shown in **Figure 1** c, with fibres curving around each other without fusing. At high magnification, the fibre surfaces exhibited pronounced dimples.

**Table 1.** Morphometric properties of the electrospun scaffold for subcutaneous and circulatory implantation

|  | **Scaffold for subcutaneous implantation** | **Scaffold for circulatory implantation** |
| --- | --- | --- |
| Wall thickness (mm) | $1.5 \pm 0.5$ | $0.9 \pm 0.4$ |
| Porosity (%) | $85.4 \pm 2.2$ | $82.0 \pm 3.5$ |
| Fibre diameter ($\mu\text{m}$) | $6.7 \pm 2.4$ | $10.5 \pm 4.7$ |
| Dispersion ($^{\circ}$) | $12.4 \pm 4.0$ | $23.2 \pm 9.5$ |

**Figure 1.**
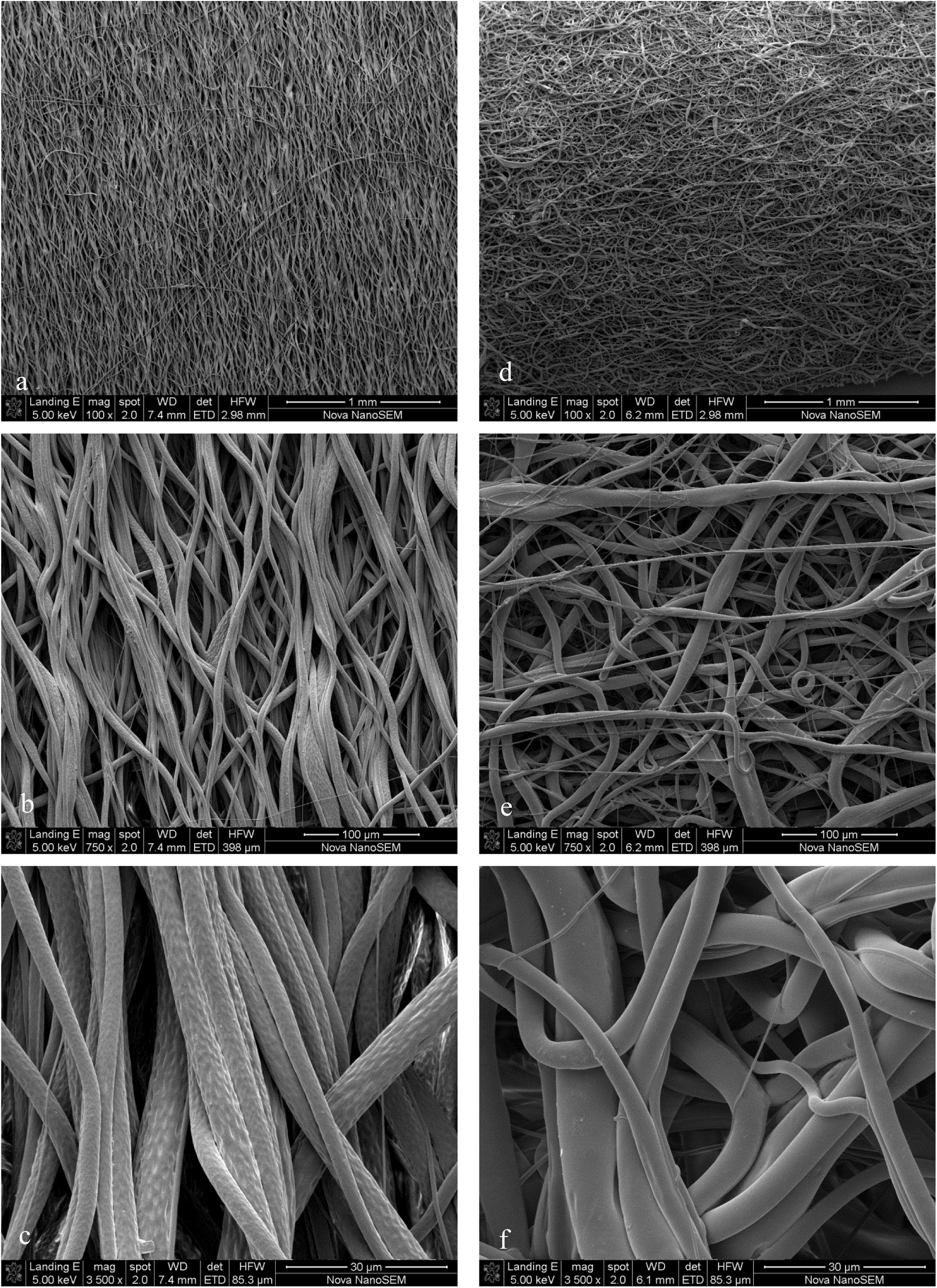
SEM micrographs of fibrous scaffold for subcutaneous (a-c) and circulatory (d-f) implantation at magnifications of 100x (a, d), 750x (b, e), and 3500x (c, f).

The scaffold for circulatory implantation exhibited fibres with considerable variation in thickness and random orientation. The low goodness factor of 0.72 of the fibre dispersion of 23 ± 9% (180° corresponds to a random distribution) indicated limited accuracy of the Gaussian fit for dispersion and suggested that fibres were more dispersed than the measured 23°. Similar to the subcutaneous scaffold, little fibre merging was observed, however the fibre surface was much smoother with substantial less dimpling.

### 3.2 Tissue ingrowth

Histological micrographs illustrating the ingrowth of tissue in the scaffold after 7, 14 and 28 days are provided in **Figure 2** for subcutaneous implants and in **Figure 3** for circulatory implants. Tissue ingrowth after 7, 14 and 28 days was 65.0 ± 13.0%, 87.3 ± 3.9% (p = 0.008) and 96.5 ± 2.3% (p = 0.001), respectively, for the subcutaneous position and 64.9 ± 9.8%, 74.3 ± 9.9% (not significant, ns) and 77.8 ± 5.4% (ns), respectively, for the aortic interposition (P values refer to comparison to t = 7 days). A decrease in gauge length of 22% was observed for subcutaneous samples implanted for 28 days compared to those implanted for 14 days despite a consistent pre-implant sample length for the entire study (18 ± 0.5 mm). A moderate and dispersed deposition of collagen could be observed at 14 and 28 days (**Figure 2** f and i, respectively)

**Figure 2.**
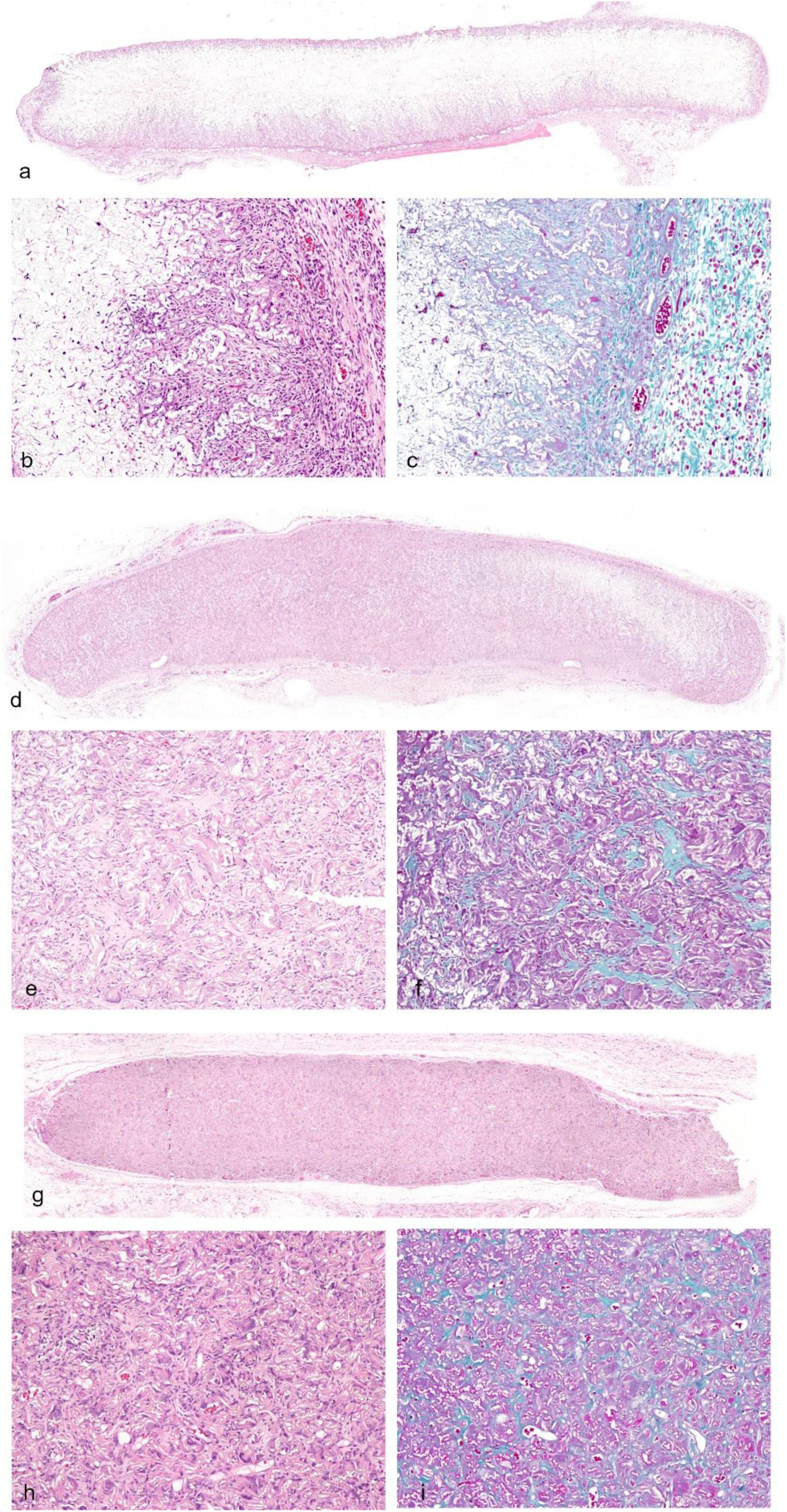
Histology micrographs of H&E and Masson’s Trichrome sections from mid region of scaffold samples after 7 days (a-c), 14 days (d-f), and 28 days (g-i) of subcutaneous implantation.

**Figure 3.**
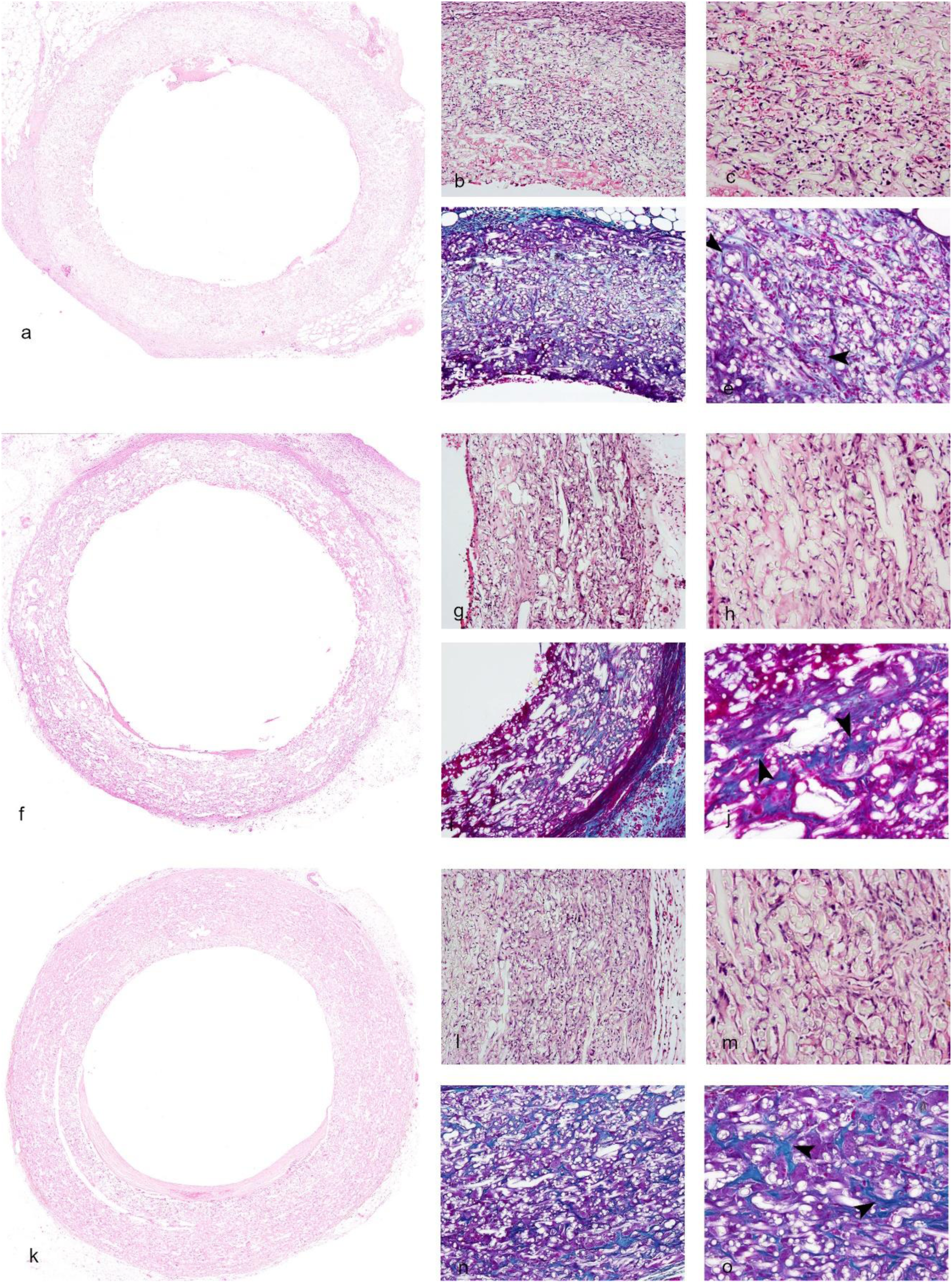
Histology micrographs of H&E and Masson’s Trichrome sections from the mid region of the scaffold samples after 7 days (a-e), 14 days (f-j), and 28 days (k-o) of circulatory implantation. Arrow heads in the 20x Masson’s Trichrome micrographs (e, j, o) identify collagen observed at all three time points.

Cellular invasion can be seen to occur from the adventitial side towards the lumen of circulatory implants at 7 days (**Figure 3** a-c) with an acellular band (possibly fibrin) bordering the lumen. Also erythrocytes are visible within the wall of the graft at 7 days suggestive of low level insudation from the circulation. More complete cellular occupation of the wall can be observed at 14 days (**Figure 3** f-h) relative to 7 days becoming more dense at 28 days (**Figure 3** k, m). Vessels were present within the wall from 14 days onwards (supplementary Figures 1-2). Even at 28 days, no evidence of a chronic inflammatory response to the scaffold in the form of foreign body giant cells could be found. A low level of collagen with a scattered deposition was found at 7 days (**Figure 3** d, e) with a similar pattern of collagen staining could be seen at the later time points that moderately increased in density.

### 3.3 Mechanical properties

**Figure 4** (a) provides graphs of stress versus strain for the subcutaneous scaffold before implantation (0 day) and after 7, 14, and 28 days implant duration in circumferential and axial direction of the scaffold for a maximum strain of 25%. In circumferential direction, little change was observed from 0 to 7 and 14 days implant duration whereas there was a substantial decrease in stress after 28 days implantation. In axial scaffold direction, an increase of stress from pre-implant scaffold and implanted scaffolds was observed. **Figure 4** (b) illustrates stress versus strain of up to 17% in circumferential direction of the circulatory graft samples at 0, 7, 14, and 28 days implantation. For these samples, a significant increase in stress was observed from 0 to 7 days, whereas changes were smaller thereafter.

**Figure 4.**
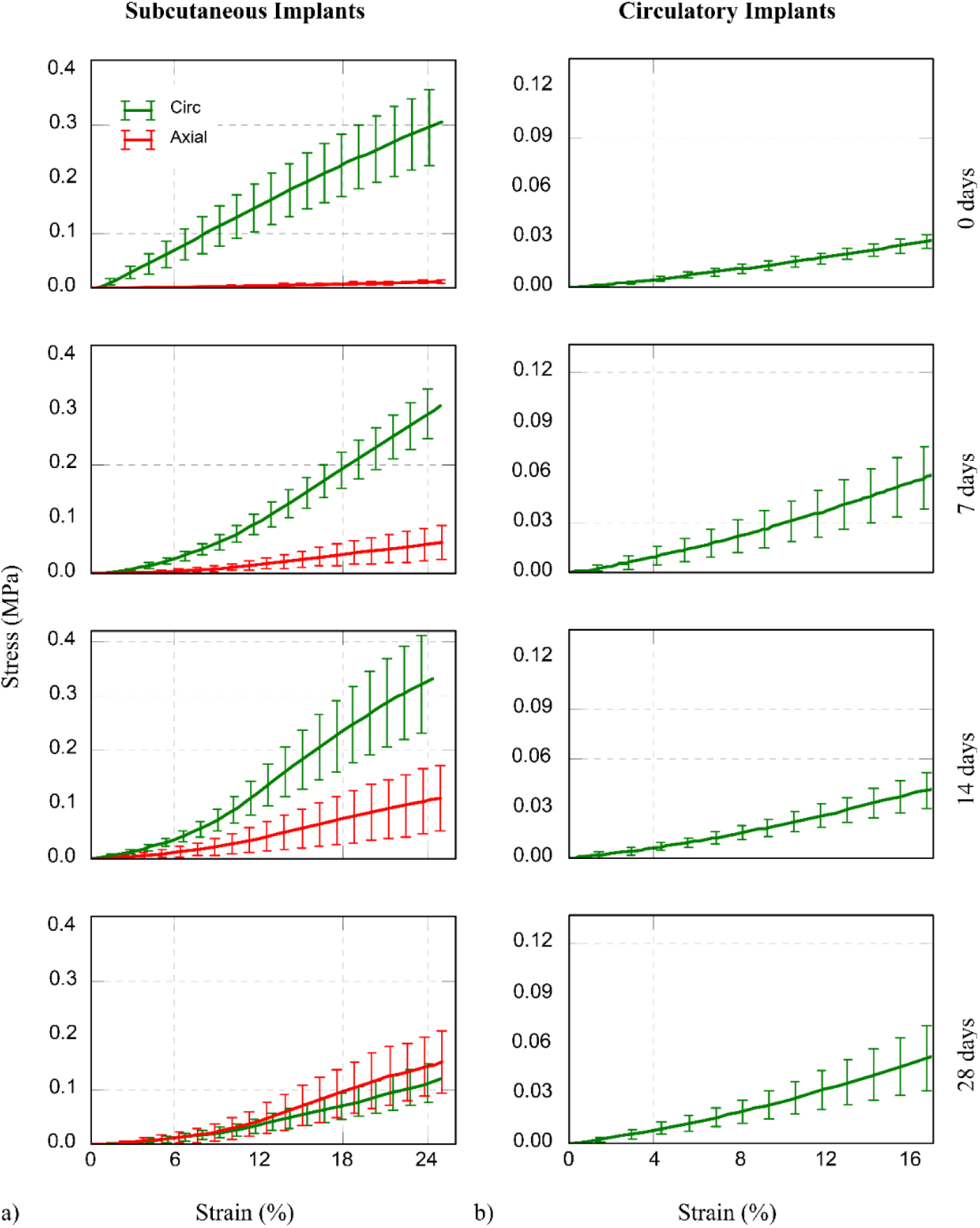
Stress versus strain curves for scaffolds before implantation (0 day) and after implantation for 7, 14 and 28 days for (a) circumferential and axial direction of scaffolds of subcutaneous implants, and (b) circumferential direction of scaffolds of circulatory implants.

**Figure 5** shows stress at 12% and 16% strain and elastic modulus at 6% and 12% strain versus implant time in circumferential and axial scaffold directions for the subcutaneous implants (**Figure 5** a, b) and in circumferential direction only for the circulatory implants (**Figure 5** c, d). In circumferential direction of the subcutaneous scaffold implants, stress was significantly lower after 28 days implantation compared to the 0 day pre-implant (p < 0.0005). The circulatory scaffold implants exhibited an increase of the elastic modulus at 6% strain by 147% (p < 0.005) and at 12% strain by 77% (ns) between T = 0 and 28 days (**Figure 5** d).

**Figure 5.**
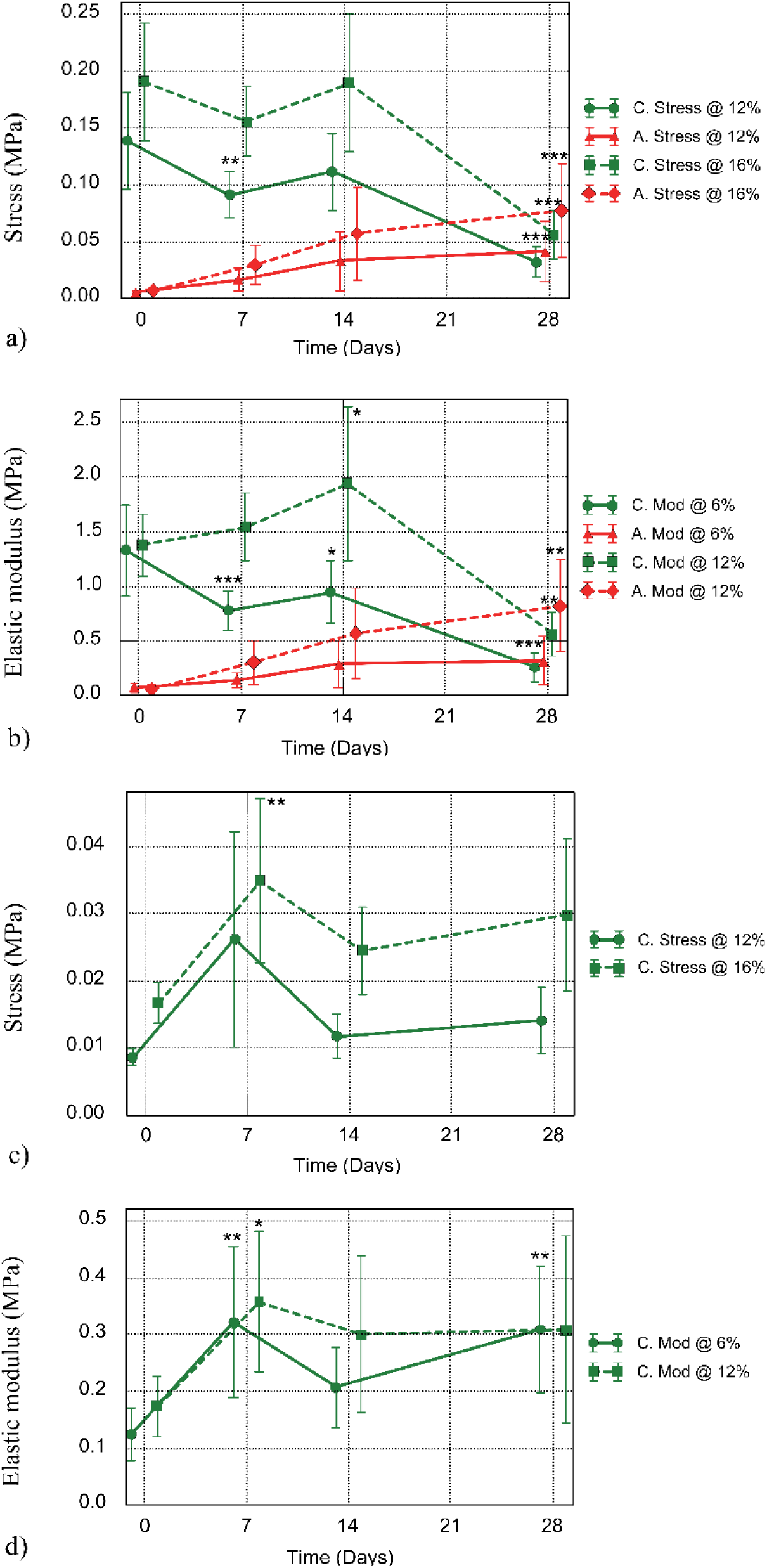
a) Stress at a strain of 12% and 16% versus implantation time and (b) elastic modulus at a strain of 6% and 12% versus implantation time for the circumferential (C) and axial (A) direction of the scaffold of subcutaneous implants. (c) Stress at a strain of 12% and 16% versus implantation time and (d) elastic modulus at the strain of 6% and 12% versus implantation time for the circumferential direction of the scaffold of circulatory implants. (*, ** and *** indicate p < 0.05, 0.005 and 0.0005, respectively, for comparison to t = 0 days)

## 4. Discussion

Tissue ingrowth into scaffolds was similar for the subcutaneous implants and the circulatory implants during the first 7 days. Thereafter, ingrowth was slower in the circulatory compared to the subcutaneous implants.

The difference in fibre thickness between the two scaffolds was most likely attributed to the different spinning target and rotational speed, as the uptake speed affects fibre thickness.^16^ The fibres of the circulatory scaffold were smooth and did not exhibit the dimpled surface texture that was present in the subcutaneous scaffolds. With comparable porosities, these two aspects could have contributed to deeper tissue penetration in the subcutaneous implants compared to the circulatory implants.

However, the different characteristics of the implant sites were the more likely cause, as subcutaneous samples were not immediately exposed to mechanical loading and were not clotted with blood at the time of implant.

Morphological differences of the scaffolds were observed; the scaffolds for assessment in subcutaneous positions exhibited a substantially higher degree of fibre alignment than the graft scaffolds used in the circulatory position. The difference was attributed to larger diameter of the electrospinning mandrel used for subcutaneous scaffolds, resulting in higher velocities of fibre uptake and consequently fibre alignment, compared to that used for the scaffolds for circulatory interposition.

### *Ex vivo* scaffold mechanics of subcutaneous implants

The decrease in circumferential stiffness between 14 to 28 days (**Figure 5** b) was ascribed to increasing morphological disruption of the highly aligned scaffold by ingrowing tissue, facilitated by the low degree of fibre merging. Displacement of fibres by ingrowing tissue is mostly expected to occur in axial scaffold direction, i.e. transverse to circumferentially aligned fibres. This proposed mechanism is illustrated in **Figure 6**. With progressing tissue ingrowth, the scaffold becomes increasingly compacted. The local realignment of the fibres during compaction leads to shortening of the scaffold in circumferential direction (as observed for samples of T = 28 days compared to T = 24 days) and an increase in structural stiffness (modulus) due to inhibition of local fibre realignment. The shortening of the scaffolds observed in the current study is slightly different from findings of van Vlimmeren et al. ^21^ for tissue engineered heart valves. They reported that tissue compaction was initiated when the biodegradable polyglycolic acid/poly-4-hydroxybutyrate scaffolds lost their mechanical integrity. As Pellethane® is biostable, the mechanical integrity of the scaffold did not change *in vivo*. The main cause for the two effects is, however, the same: ingrowing tissue exerts forces on the scaffold which modifies the scaffold geometry.

**Figure 6.**
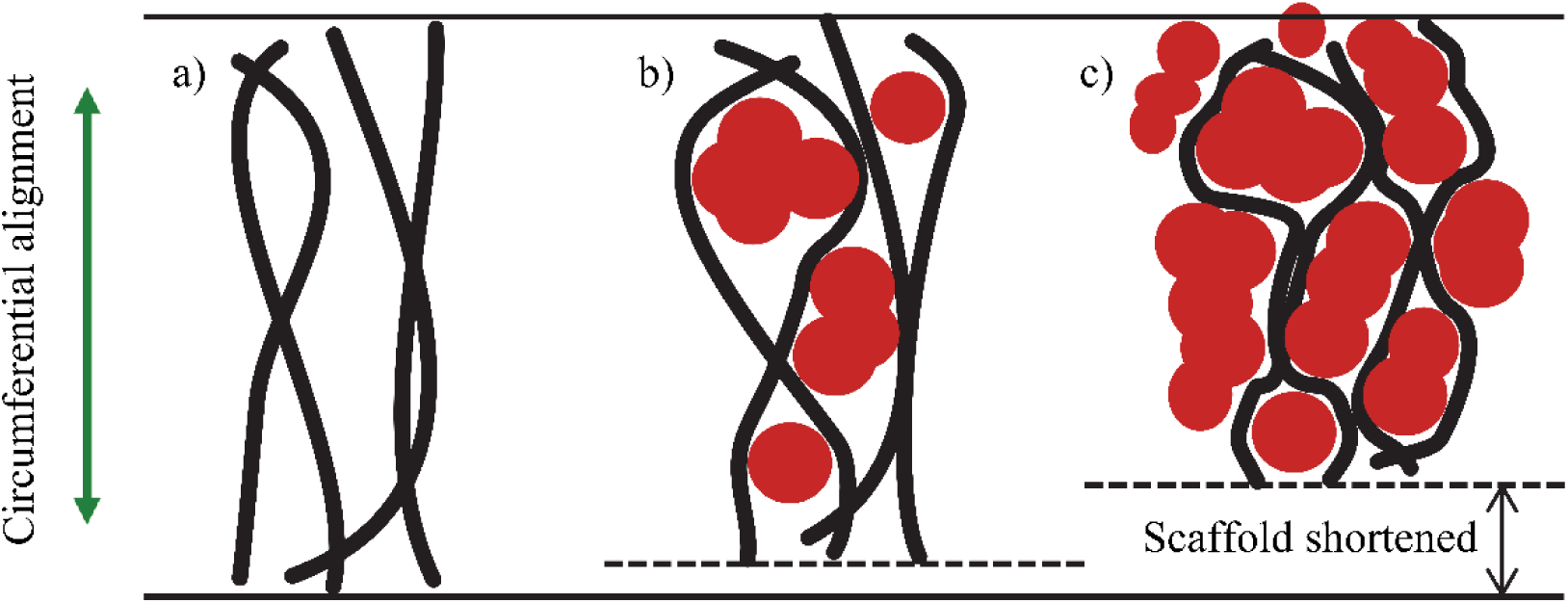
Fibre alignment changes and scaffold shortening due to tissue ingrowth. Fibres (black) of a pre-implant scaffold (a) are displaced by ingrowing tissue (red, b and c). During early ingrowth (b), the scaffold structure still allows local re-alignment of fibres. Progressing tissue ingrowth (c) leads to increasing compaction of the scaffold and space for local realignment of fibres decreases.

*Increase in axial stiffness with increasing implant duration (***Figure 5** *b):* A significant increase of the elastic modulus at 12% strain in axial scaffold direction between T = 0 and T = 28 days (p < 0.005) is based on the low elastic modulus in the scaffold pre-implantation transverse to the predominant fibre alignment, and the large mechanical contribution the ingrowing tissue provides relative to the low pre-implant axial scaffold stiffness. The continuous increase in elastic modulus, although not statistically significant at 7 and 14 days, is ascribed to the progressive ingrowth of tissue observed.

*Change in stress-strain behaviour with increasing implant duration:* There was a change in shape of the stress-strain curves with the incorporation of tissue in the scaffold. At T = 0 days, the stress-strain curve is fairly linear, based only on the mechanical and structural properties of the fibrous scaffold. From T=7 days onwards, the curves indicate non-linear stiffening in the 0-12% strain range (**Figure 4** a), as is generally associated with tissue.^8, 11, 23, 27^ The change from linear to non-linear elastic behaviour is reflected in a decrease in the elastic modulus at 6% strain of 41% and 29% and an increase in the elastic modulus at 12% strain of 11% and 41% from T = 0 days to 7 days and 14 days, respectively (**Figure 5** b). The change in the stress-strain response is attributed to the increasing contribution of ingrowing tissue to the mechanical behaviour of the scaffold, and the associated local changes in fibre alignment already discussed. Local displacement and de-alignment of fibres by newly formed tissue (**Figure 6** b) leads to low stiffness associated with the wavy state of the fibres and higher stiffness once the fibres are straightened under increasing extension of the scaffold. The largest change in the elastic modulus compared to pre-implant time point was observed in the circumferential scaffold direction at 6% strain after 28 days of implantation.

### *Ex vivo* scaffold mechanics of circulatory implants

An unexpected observation for the circulatory implants was that these scaffolds exhibited the maximum elastic modulus after 7 days of implantation. For the subcutaneous and the circulatory studies, samples retrieved at T = 7 days displayed more surrounding tissue than at 14 and 28 days. Removing this tissue capsule without damaging the scaffold was more intricate for the vascular graft samples than for the flat samples. Remaining excess tissue is the most likely explanation for our finding. Disregarding the T = 7 days time point, the stress at 12% and 16% strain (**Figure 5** c) and the elastic modulus at 6% strain increased gradually until 28 days implantation, whereas the elastic modulus at 12% strain increased until T = 14 days but did not change thereafter. Although not statistically significant, the elastic modulus increased with increasing strain from 6% to 12% by 39%, 11% and 45% at T = 0, 7 and 14 days, respectively, but did not increase at T = 28 days.

### Limitations of the study

The difference in scaffold morphology limited the direct comparison of results from the subcutaneous and the circulatory implant position. However, different sample geometries for the two implant positions required differences in the electrospinning process parameters e.g. target geometry and associated angular velocity of the surface of the rotating target. While optimisation of the process parameters may provide more similar scaffold morphology, the comparison of the implant positions in the current study was of exploratory nature.

Whereas the mechanical properties of the scaffold may still have changed beyond 28 days of implantation, the current study was limited to 28 days in the context of fast degrading scaffolds that we studied *in vitro* previously.^12^ We observed that elastic modulus and mechanical strength of the biodegradable scaffold decreased predominantly during the first 14 days of hydrolytic degradation, but less thereafter.

The measurement of mechanical properties in the axial direction of circulatory implanted scaffolds would have provided additional data. However, the predominant random fibre orientation of the circulatory samples resulted in a low degree of mechanical anisotropy and similar mechanical properties of the scaffold in different directions. Hence, mechanical properties of the circulatory implanted scaffold in axial and circumferential directions do not differ to the same degree as for the subcutaneous scaffold. In combination with the small sample dimensions of circulatory implants and single implant position per animal, the limited benefit of axial measurements did not justify doubling the number of animals involved in the circulatory sub-study.

Considering the small pre-implantation wall thickness of the scaffolds (1.5 ± 0.5 mm for subcutaneous implants and 0.9 ± 0.4 mm for circulatory implants), the measurement of wall thickness after sample retrieval did not provide data suitable to assess the change in wall thickness due to scaffold compaction. In addition, length changes due to scaffold compaction was observed predominantly in direction of the fibre alignment, but not transverse to the fibre alignment, of the highly aligned scaffolds for subcutaneous implants. Changes in the wall thickness of the samples would as such be difficult to discern.

## Conclusions

The softening of the highly aligned scaffold observed in the predominant fibre direction during formation of tissue poses a challenge in scaffold design. The scaffold softening not only limits the stiffness and strength of tissue engineered constructs; it also attenuates the mechanical anisotropy that may be desired for functional purposes. These effects may alter the desired long-term performance of the implants and need to be further studied.

## Acknowledgements

This study was supported financially by the National Research Foundation (NRF) of South Africa and the South African Medical Research Council (SAMRC). Any opinion, findings and conclusions or recommendations expressed in this publication are those of the authors and therefore the NRF and SAMRC do not accept any liability in regard thereto. HK received a matching dissertation grant from the International Society of Biomechanics. JK acknowledges financial support by Deutsche Herzstiftung (Jahresstipendium S/08/10). The authors thank Helen Ilsley of the Cardiovascular Research Unit, University of Cape Town, for the excellent technical assistance with histological sections.

## Data

Raw data for fibre diameter, fibre alignment, tissue ingrowth and mechanical characterisation is available on ZivaHUB (http://doi.org/10.25375/uct.9888710).

