## Supplementary figures and images for "Tissue ingrowth markedly reduces mechanical anisotropy and stiffness in fibre direction of highly aligned electrospun polyurethane scaffolds"

### Supplemental Figure 1

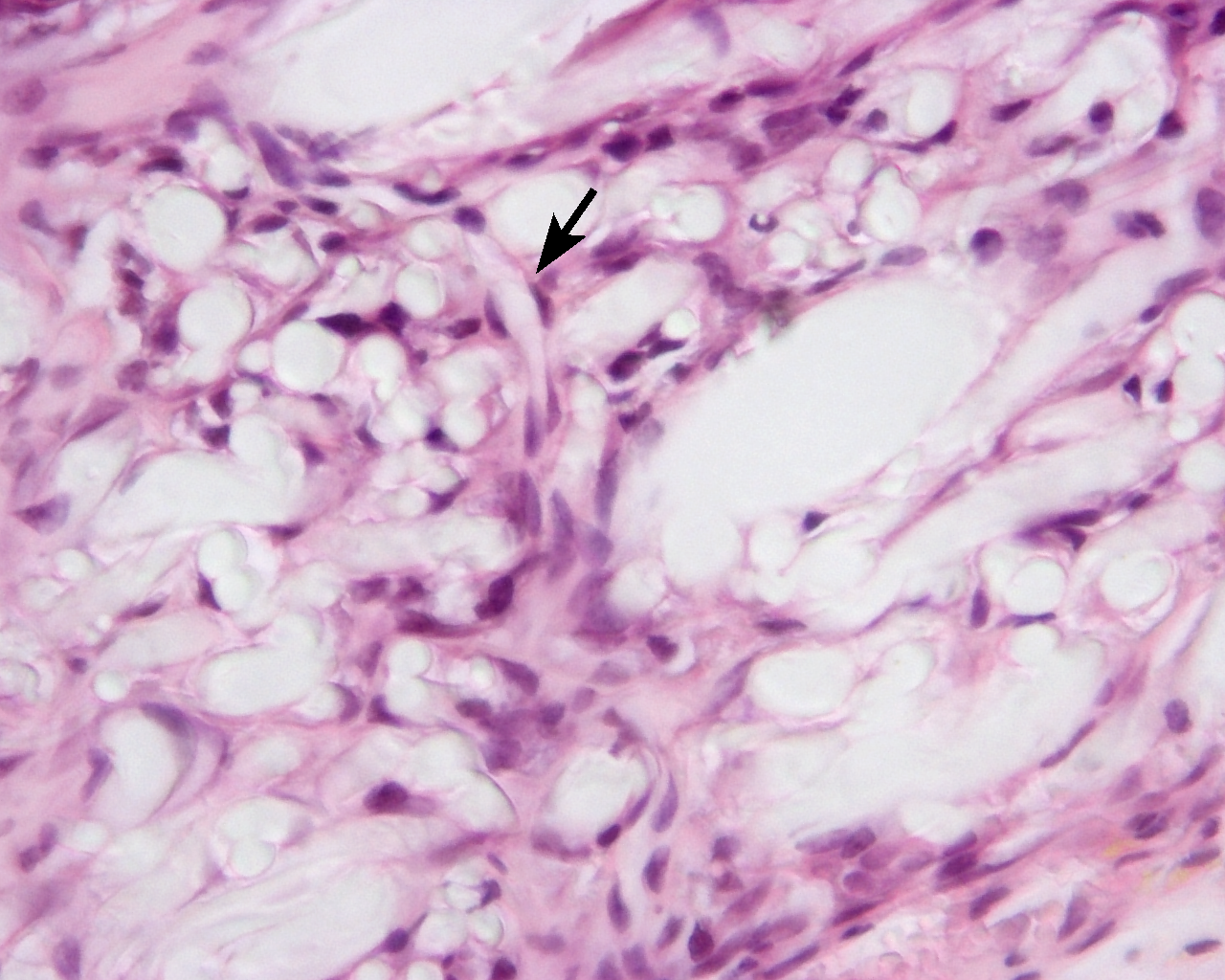

### Supplemental Figure 2

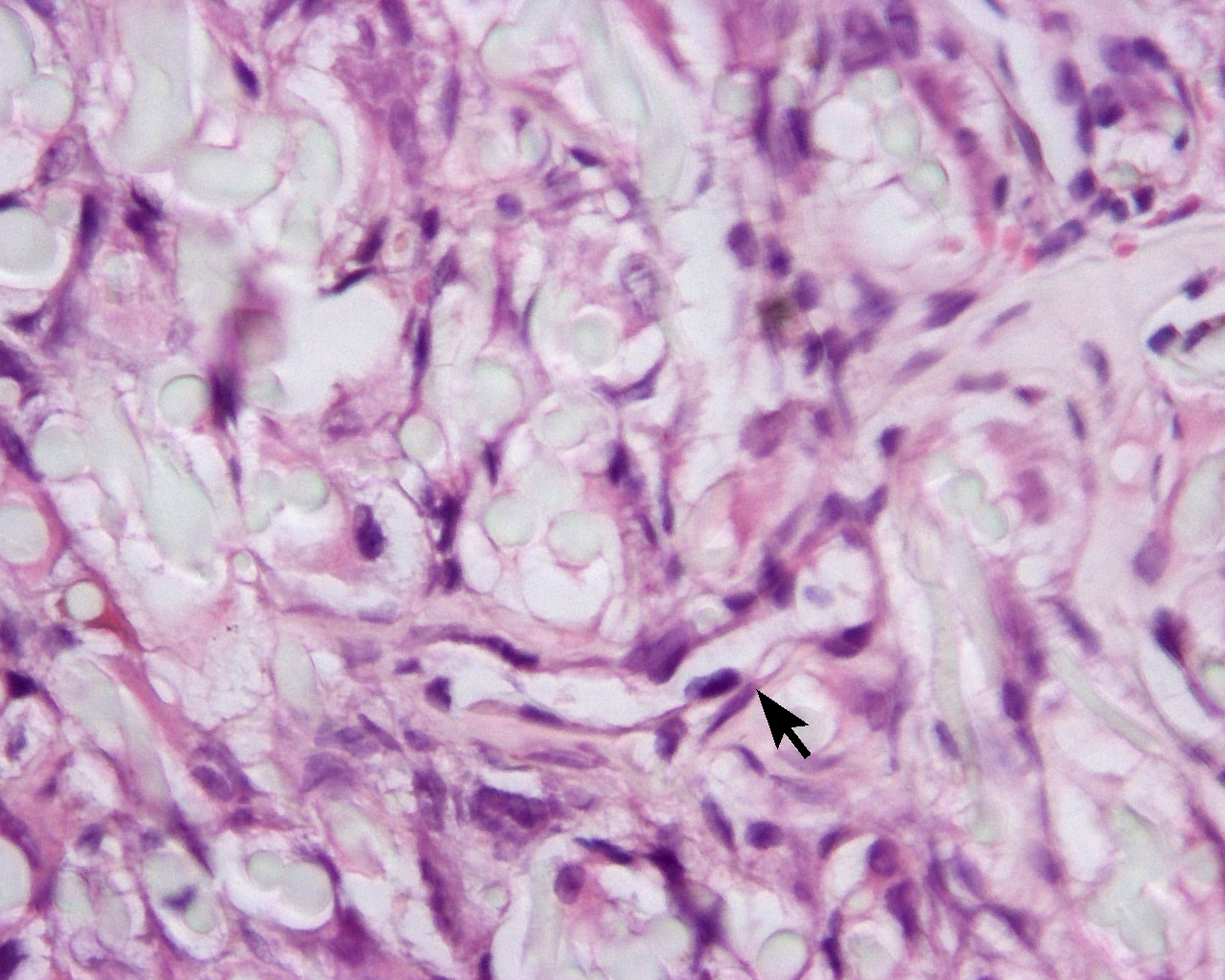
